# Functional hierarchy among different Rab27 effectors involved in secretory granule exocytosis

**DOI:** 10.1101/2022.08.23.504970

**Authors:** Kunli Zhao, Kohichi Matsunaga, Kouichi Mizuno, Hao Wang, Katsuhide Okunishi, Tetsuro Izumi

## Abstract

The Rab27 effectors play versatile roles in regulated exocytosis. In pancreatic beta cells, exophilin-8 anchors granules in the peripheral actin cortex, whereas granuphilin and melanophilin mediate granule fusion with and without stable docking to the plasma membrane, respectively. However, it is unknown whether these coexisting effectors function in parallel or in sequence to support the whole insulin secretory process. Here, we investigate their functional relationship by comparing the exocytic phenotypes in beta cells simultaneously lacking two effectors with those lacking one of them. Analyses of prefusion profiles by total internal reflection fluorescence microscopy indicate that melanophilin exclusively functions downstream of exophilin-8 to mobilize granules for fusion from the actin network to the plasma membrane after stimulation. The two effectors are physically linked via the exocyst complex. Downregulation of the exocyst component affects granule exocytosis only in the presence of exophilin-8. Exophilin-8 also promotes fusion of granules stably docked to the plasma membrane mediated by granuphilin, although it is dispensable for granule residence beneath the plasma membrane. The current study presents the first diagram for multiple intracellular paths of granule exocytosis and for functional hierarchy among different Rab27 effectors in the same cell.

## Introduction

Synaptic vesicles should have been docked and primed on the plasma membrane before stimulation to execute neuronal transmission within 1 millisecond after electrical stimulation (Südhof, 2013). In contrast, secretory granules in exocrine and endocrine cells initiate exocytosis at a more than thousand times later time point even if Ca^2+^ concentration is abruptly upregulated by caged-Ca^2+^ compounds (Kasai, 1999). Furthermore, they physiologically release bioactive substances in much more prolonged time: for example, digestive enzymes and insulin are secreted from pancreas continuously in a range of minutes and sometimes even hours during food intake and subsequent hyperglycemia. To support such slow and consecutive exocytosis, the exocytic pathway may not be single or linear, and steps to recruit granules from the cell interior to the cell limits is thought to be critical. In fact, total internal reflection fluorescence (TIRF) microscopy that can monitor prefusion behavior in living cells reveals that insulin granules residing beneath the plasma membrane before stimulation and those recruited from the cell interior after stimulation fuse in parallel, though their ratios may change, during a physiological time course of insulin secretion (Kasai et al., 2008; Ohara-Imaizumi et al., 2004; Shibasaki et al., 2007). In many secretory cells, granules are clustered in the actin cortex at the cell periphery and/or along the plasma membrane, compared with other cytoplasmic areas. It has traditionally been considered that granules docked to the plasma membrane form a readily releasable pool, whereas those accumulated within the actin cortex form a reserve pool. However, this scenario may be too simplified, considering that there is a gating system to prevent spontaneous or unlimited vesicle fusion in regulated exocytosis. In fact, multiple Rab27 effectors that are involved in intracellular granule trafficking show complex and differential effects on exocytosis (Izumi, 2021). For example, granuphilin (also known as exophilin-2 and Slp4) mediates stable granule docking to the plasma membrane but simultaneously prevents their spontaneous fusion by interacting with a fusion-incompetent, closed form of syntaxins (Gomi et al., 2005; Torii et al., 2002). Another effector, exophilin-8 (also known as MyRIP and Slac2-c), anchors secretory granules within the actin cortex (Bierings et al., 2012; Desnos et al., 2003; Fan et al., 2017; Huet et al., 2012; Mizuno et al., 2011; Nightingale et al., 2009), which is considered to have dual roles in accumulating granules at the cell periphery and in preventing their access to the plasma membrane. Although another effector, melanophilin (also known as exophilin-3 and Slac2-a), similarly captures melanosomes within the peripheral actin network in skin melanocytes (Hammer and Sellers, 2012), it mediates stimulus-induced granule mobilization and immediate fusion to the plasma membrane in pancreatic beta cells (Wang et al., 2020). However, it remains unknown how different Rab27 effectors coexisting in the same cell function in a coordinated manner to support the whole exocytic processes. It has not been established either whether multiple secretory pathways and/or rate-limiting steps exist toward the final fusion in regulated granule exocytosis. To explore these problems, we must first understand whether each effector functions in sequence or in parallel. In the present study, we compared the exocytic profiles in beta cells lacking the two effectors with those in cells deficient in each single effector and in wild-type (WT) cells. As a result, we obtained the information about the functional hierarchy and relationship among the exocytic steps in which individual effectors are involved. Furthermore, we found that the exocyst, which universally functions in constitutive exocytosis (Wu and Guo, 2015), is involved in physical and functional connection between different Rab27 effectors.

## Results

### Melanophilin exclusively functions downstream of exophilin-8 to mediate the exocytosis of granules recruited from the actin cortex to the plasma membrane after stimulation

In monolayer mouse pancreatic beta cells, insulin granules were unevenly distributed with accumulation in the actin cortex (Figure 1—figure supplement 1A). Because the Rab27 effectors, melanophilin and exophilin-8, are known to show affinities to actin motors, myosin-Va and/or - VIIa (Hammer and Sellers, 2012; Izumi, 2021), these effectors are expected to function on granules within this peripheral actin network. In fact, they showed a similar uneven distribution and were colocalized especially at the cell periphery (Figure 1—figure supplement 1B). To examine the functional relationship between these effectors, we generated melanophilin/exophilin-8 double-knockout (ME8DKO) mice by crossing exophilin-8-knockout (Exo8KO) mice (Fan et al., 2017) with melanophilin-knockout (MlphKO) mice (see Materials and methods). The doubly deficient beta cells displayed even distribution of insulin granules in the cytoplasm without accumulation at the cell periphery, in contrast to WT and MlphKO cells, but at similarity to Exo8KO cells (Figure 1A). Expression of exophilin-8, but not that of melanophilin, in ME8DKO cells at the endogenous level in WT cells redistributed them to the cell periphery (Figure 1B). These findings confirm the previous findings obtained from cells deficient in each single effector that exophilin-8 is essential for granule accumulation within the actin cortex, whereas melanophilin is not (Fan et al., 2017; Wang et al., 2020).

**Figure 1.**
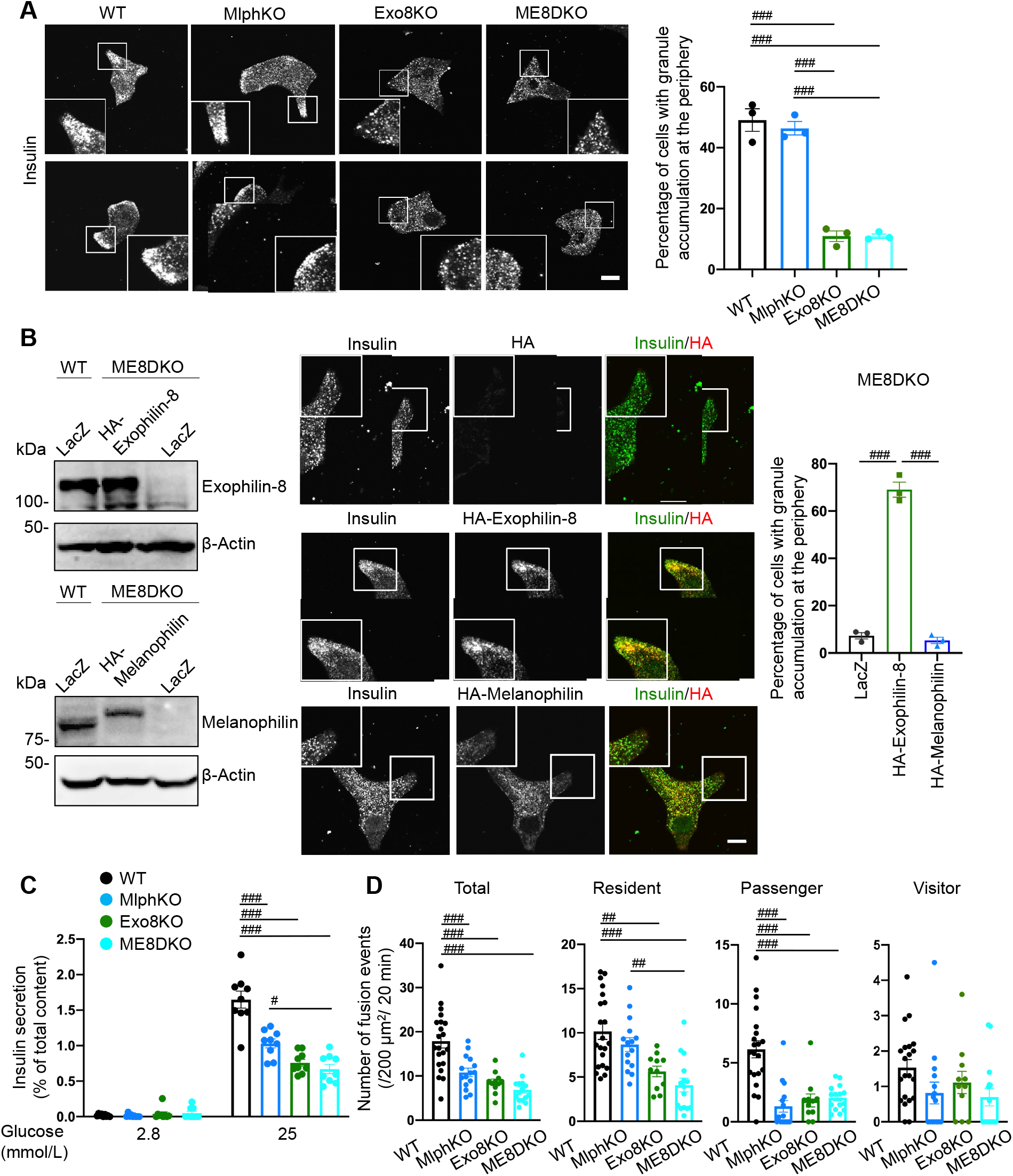
Insulin secretory defects of beta cells doubly deficient in melanophilin and exophilin-8 are indistinguishable from those singly deficient in exophilin-8. **A**: WT, MlphKO, Exo8KO, and ME8DKO beta cells were immunostained with anti-insulin antibody. A peripheral accumulation of insulin immunosignals was quantified under confocal microscopy: clustering of insulin immunosignals at least in one corner (upper) or along the plasma membrane (lower) were counted as positive. More than 100 cells were inspected in each of three independent experiments. Note that Exo8KO and ME8DKO cells do not show the peripheral accumulation of insulin granules, in contrast to WT and MlphKO cells. **B:** ME8DKO cells were infected by adenovirus expressing either HA-exophilin-8 or HA-melanophilin at the endogenous protein levels found in WT cells. LacZ was expressed in WT and ME8DKO cells as controls. The cell extracts were immunoblotted with the indicated antibodies (left). The ME8DKO cells expressing LacZ (upper), HA-exophilin-8 (middle), and HA-melanophilin (lower) were immunostained with anti-insulin and anti-HA antibodies (center), and a peripheral accumulation of insulin was quantified as in **A** (right). Insets represent higher magnification photomicrographs of a cell within the region outlined by frames. Note that expression of HA-exophilin-8, but not HA-melanophilin, rescues the peripheral granule accumulation in ME8DKO cells. **C**: Islets isolated from WT, MlphKO, Exo8KO, and ME8DKO mice at 12-18 weeks of age were preincubated in 2.8 mmol/L low glucose (LG)-containing KRB buffer at 37°C for 1 h. They were then incubated in new LG buffer for 1 h followed by 25 mmol/L high glucose (HG) buffer for 1 h. Insulin levels secreted in the media and left in the cell lysates were measured, and their ratios are shown (*n* = 9 from 3 mice each). **D**: A monolayer of islet cells from WT (*n* = 21 cells from 5 mice), MlphKO (*n* = 10 cells from 3 mice), Exo8KO (*n* = 12 cells from 3 mice), and ME8DKO (*n* = 12 cells from 3 mice) mice were infected by adenovirus encoding insulin-EGFP. Insulin granule fusion events in response to 25 mmol/L glucose for 20 min were counted under TIRF microscopy. Fusion events observed were categorized into three types, residents, visitors, and passengers. Note that the decrease in insulin exocytosis in ME8DKO cells is severer than that in MlphKO cells, specifically due to the decrease in the resident type exocytosis. Bars, 10 μm. ## *P* < 0.01, ### *P* < 0.001 by one-way ANOVA. **Source data 1.** Uncropped blot images of Figure 1B.

The cells deficient in melanophilin and/or exophilin-8 all showed decreases in glucose-stimulated insulin secretion (GSIS) compared with WT cells, although the decrease in ME8DKO cells was larger than that in MlphKO cells (Figure 1C). We next monitored insulin granule exocytosis directly by TIRF microscopy in living cells expressing insulin fused with enhanced green fluorescent protein (EGFP). We previously categorized fused insulin granules into three types depending on their prefusion behaviors (Kasai et al., 2008): those having been visible before stimulation, ‘residents’; those becoming visible during stimulation, ‘visitors’; and those invisible until fusion, ‘passengers’. In good correlations with the amount of insulin released in the medium, the total number of fusion events detected as a flash followed by diffusion of insulin-EGFP fluorescence during glucose stimulation was decreased in each of the mutant cells (Figure 1D). MlphKO cells with the genetic background of C57BL6/N mice showed a specific decrease in the passenger type exocytosis compared with WT cells, as found in parental *leaden* cells with the genetic background of C57BR/cdJ mice (Wang et al., 2020). In contrast, both Exo8KO and ME8DKO cells with the same C57BL6/N genetic background displayed decreases in both resident and passenger types of exocytosis, although a much less frequent, visitor type was difficult to compare among the cells. Because there are no significant differences in exocytic phenotypes between Exo8KO and ME8DKO cells, melanophilin is thought to function downstream of exophilin-8. Considering the previously identified role of each effector in beta cells (Fan et al., 2017; Wang et al., 2020), the passenger type exocytosis mediated by melanophilin via interactions with myosin-Va and syntaxin-4 appears to be derived from granules anchored in the actin cortex by exophilin-8.

### Exophilin-8 and melanophilin form a complex via the exocyst

To explore the molecular basis that exophilin-8 and melanophilin sequentially promote the passenger type exocytosis, we first examined the expression level of each effector in cells lacking the other effector. Although the expression of exophilin-8 was not affected in MlphKO cells, that of melanophilin was decreased by half in Exo8KO cells (Figure 2A). We further found that exophilin-8 and melanophilin expressed in MIN6 cells form a complex (Figure 2B). These findings suggest that the protein stability of melanophilin partially depends on its interaction with exophilin-8, which is consistent with the model that melanophilin functions downstream of exophilin-8. However, they do not interact when expressed in HEK293A cells (Figure 2—figure supplement 1A), suggesting that they do not interact directly. The two effectors have been shown to interact with different proteins, except for Rab27a, in beta cells (Fan et al., 2017; Wang et al., 2020). Exophilin-8 mutant that loses the binding activity to RIM-BP2, and melanophilin mutants that lose the binding activity to Rab27a, myosin-Va, or actin, all preserved the binding activity to the other WT effector in MIN6 cells (Figure 2—figure supplement 1B). To identify unknown intermediate proteins, we individually expressed Myc-TEV-FLAG (MEF)-tagged exophilin-8 and melanophilin in MIN6 cells, and the proteins in each of the anti-FLAG immunoprecipitates were analyzed by a liquid chromatography-tandem mass spectrometry (LC-MS/MS) system (Figure 2—figure supplement 2). As a result, we identified the exocyst complex components, SEC8, SEC10, and EXO70, in both immunoprecipitates, which is consistent with the previous finding that exophilin-8 interacts with SEC6 and SEC8 in INS-1 832/13 cells (Goehring et al., 2007). We confirmed the endogenous interactions among melanophilin, exophilin-8, SEC6, and SEC10 in MIN6 cells (Figure 2C). The exocyst forms an evolutionarily conserved heterooctameric protein complex (Wu and Guo, 2015). To identify responsible components for the interaction with two Rab27 effectors, we performed visible immunoprecipitation (VIP) assay, which can examine large numbers of protein combinations and complicated one-to-many or many-to-many protein interactions at once (Katoh et al., 2015). As a result, we found that exophilin-8 and melanophilin immediately, if not directly, interact with SEC8 and EXO70, respectively (Figure 2D). Although the exocyst components were found in both immunoprecipitates of melanophilin and exophilin-8, other interacting proteins such as RIM-BP2, RIM2, myosin-VIIa, and myosin-Va largely existed only in one of the immunoprecipitates (Figure 2E), suggesting that most of each effector forms a distinct complex in cells. In addition, although we previously reported that RIM-BP2 primarily interacts with myosin-Vlla displaying the molecular mass of ~170 kDa in INS-1 832/13 rat cells (Fan et al., 2017), it interacted with mysosin-VIIa with the authentic mass of ~260 kDa in MIN6 mouse cells, which may reflect the species difference between these beta-cell lines.

**Figure 2.**
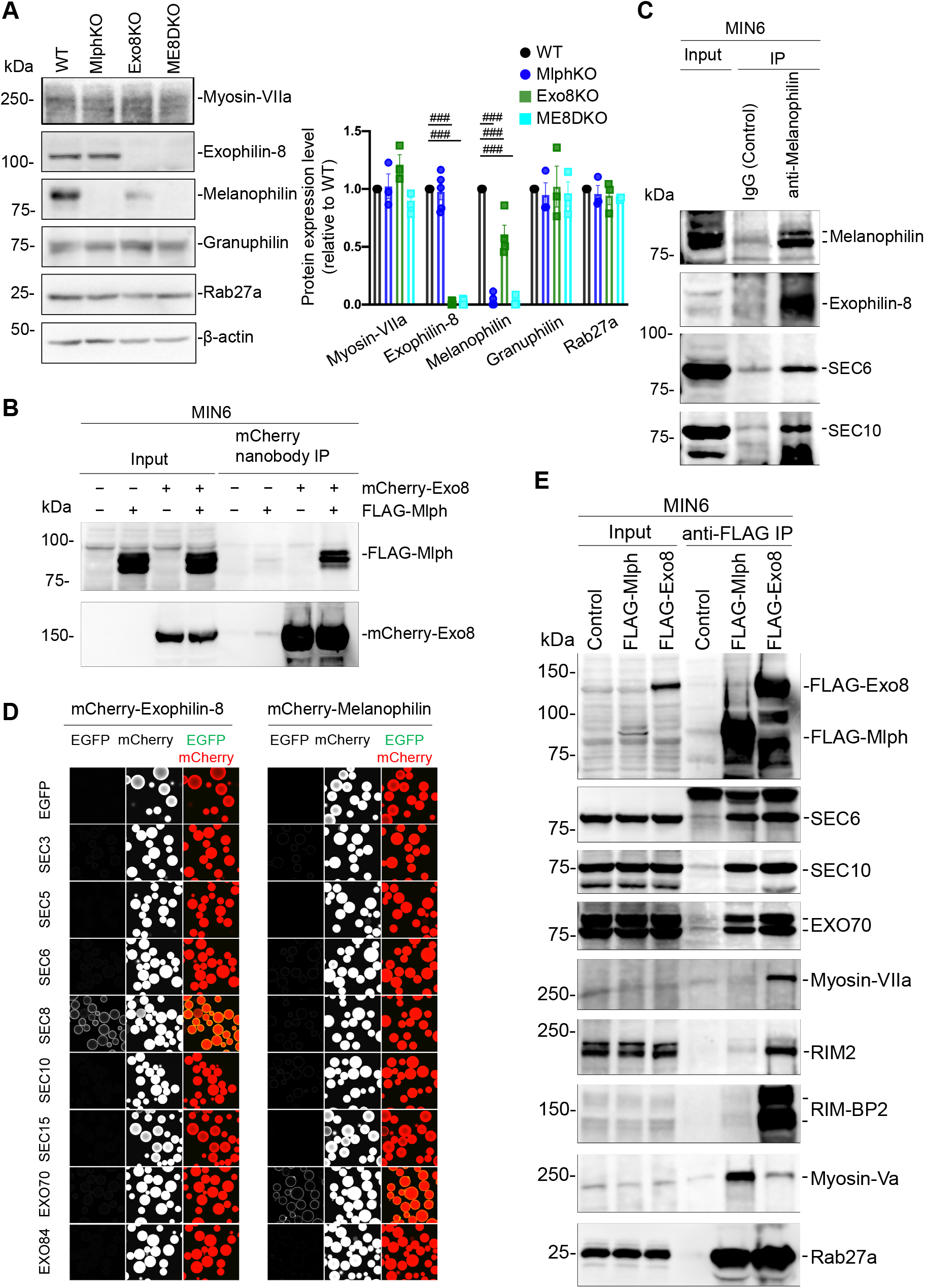
Exophilin-8 interacts with melanophilin through the exocyst in beta cells. **A**: Islet extracts (40 μg) from WT, MlphKO, Exo8KO, and ME8DKO mice were immunoblotted with antibodies toward the indicated antibodies (left). Protein expression levels were quantified by densitometric analyses from 3-5 independent preparations (right). ### *P* < 0.001 by one-way ANOVA. **B**: MIN6 cells were infected with adenoviruses encoding control LacZ, mCherry-exophilin-8 (Exo8), and/or FLAG-melanophilin (Mlph). After 2 days, the cell extracts underwent immunoprecipitation with mCherry nanobody. The immunoprecipitates (IP), as well as the 1% extracts (Input), were immunoblotted with anti-FLAG and anti-red fluorescent protein (RFP) antibodies. **C**: MIN6 cell extracts were immunoprecipitated with rabbit anti-melanophilin antibody or control IgG, and the immunoprecipitates were immunoblotted with antibodies toward the indicated proteins. **D**: HEK293A cells cultured in 10-cm dish were transfected with mCherry-fused, exophilin-8 (left) or melanophilin (right) with the indicated, EGFP-fused exocyst components. After 48 h, the cell lysates were subjected to immunoprecipitation with mCherry-nanobody bound glutathione-Sepharose beads. EGFP and mCherry fluorescence on the precipitated beads were observed by confocal microscopy. **E**: MIN6 cells expressing FLAG-tagged, melanophilin or exophlin-8 were immunoprecipitated with anti-FLAG antibody, and the immunoprecipitates were immunoblotted with antibodies toward the indicated proteins. Note that the exocyst complex components exist in both immunoprecipitates, whereas RIM-BP2, RIM2, myosin-VIIa, and myosin-Va largely exist only in one of them. **Source data 2.** Uncropped blot images of Figure 2A, B, C, and E, and figure supplement 1B.

### The exocyst functions only in the presence of exophilin-8

We next investigated the intracellular distribution of the exocyst component. As previously found in MIN6 cells (Tsuboi et al., 2005), SEC6 was colocalized with insulin granules in WT beta cells (Figure 3—figure supplement 1). It also remained on granules in MlphKO, Exo8KO, and ME8DKO cells, although its accumulation at the cell periphery was not found in exophilin-8-deficient cells due to the even granule distribution, as shown in Figure 1A. These findings indicate that the exocyst is localized on insulin granules independent of its affinities to melanophilin and/or exophilin-8. When SEC10, another exocyst component, was silenced by specific siRNAs (Figure 3—figure supplement 2A), the peripheral accumulation of insulin granules was not affected (Figure 3—figure supplement 2B), in contrast in exophilin-8-deficienct cells (Figure 1A; Figure 3—figure supplement 1). However, colocalization ratio of melanophilin, but not that of exophilin-8, with SEC6 on insulin granules was significantly decreased (Figure 3). In mammalian cells, the exocyst complex is assembled after separate formation of the subcomplex 1 (SEC3, SEC5, SEC6, SEC8) and the subcomplex 2 (SEC10, SEC15, EXO70, EXO84) (Ahmed et al., 2018). Because exophilin-8 and melanophilin immediately bind SEC8 and EXO70, respectively (Figure 2D), it is conceivable that the two effectors specifically interact with different subcomplexes. Thus, it is reasonable that silencing of SEC10 that disrupts the subcomplex 2 dissociates melanophilin from exophilin-8 associated with the subcomplex 1. These findings indicate that the exocyst physically connects the two effectors on the same granule.

**Figure 3.**
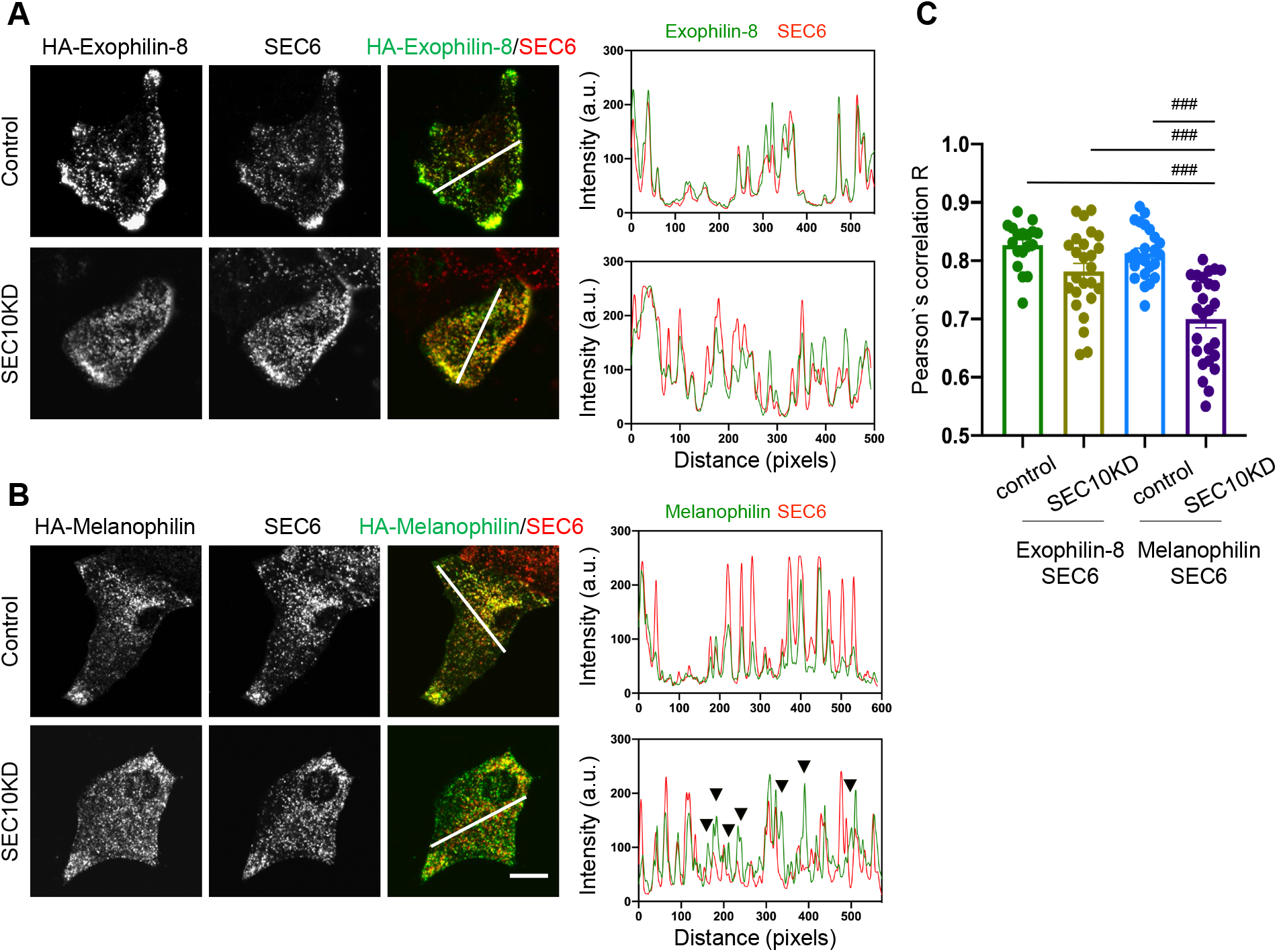
SEC10 knockdown dissociates melanophilin, but not Exophilin-8, from SEC6 on granules. HA-exophilin-8 and HA-melanophilin were expressed in Exo8KO (**A**) and MlphKO monolayer beta cells (**B**), respectively, at the endogenous levels in WT cells as described in Figure 1B. They were then transfected with control siRNA (upper) or siRNA against SEC10 #11 or #12 (lower), as shown in Figure 3—figure supplement 2A. After fixation, the cells were coimmunostained with anti-HA and anti-SEC6 antibodies and were observed by confocal microscopy (left). Fluorescent intensity profiles along the indicated line of SEC6 and either HA-exophilin-8 or HA-melanophilin are shown (right). Colocalization was quantified by Pearson’s correlation coefficient (**C**, *n* = 18-26 cells from 2 mice each). Note that SEC10 knockdown (KD) induces dissociation of melanophilin, but not exophilin-8, from SEC6 (black arrowheads). Bar, 10 μm. ### *P* < 0.001 by one-way ANOVA.

SEC10 knockdown markedly decreased GSIS in WT cells, but not further decreased GSIS in Exo8KO cells, which was already decreased by half compared with WT cells (Figure 4A). These findings suggest that the exocyst functions only in the presence of exophilin-8. TIRF microscopy of these cells expressing insulin-EGFP revealed that the number of visible granules was not decreased in Exo8KO cells (Figure 4B), which accords with the previous electron microscopic finding that the number of granules beneath the plasma membrane is only modestly decreased (Fan et al., 2017). Similarly, SEC10 knockdown does not affect the number of visible granules in either WT or Exo8KO cells. Consistent with insulin secretion assays (Figure 4A), it induces a marked decrease in glucose-stimulated fusion events in WT cells, but no additional decrease in Exo8KO cells (Figure 4C). Categorization by prefusion behavior revealed decreases in both resident and passenger types of exocytosis in WT cells, which phenocopied Exo8KO cells without SEC10 knockdown (Figure 1D). Again, SEC10 knockdown had no effects on either type of exocytosis in Exo8KO cells. The holo-exocyst appears to function with exophilin-8 in the same pathway, because SEC10 knockdown is expected to preserve the interaction between exophilin-8 and the subcomplex 1, as found in Figure 3A. Taken together, it seems that exophilin-8 and the exocyst mediate the passenger type exocytosis when they interact with melanophilin on the same granule.

**Figure 4.**
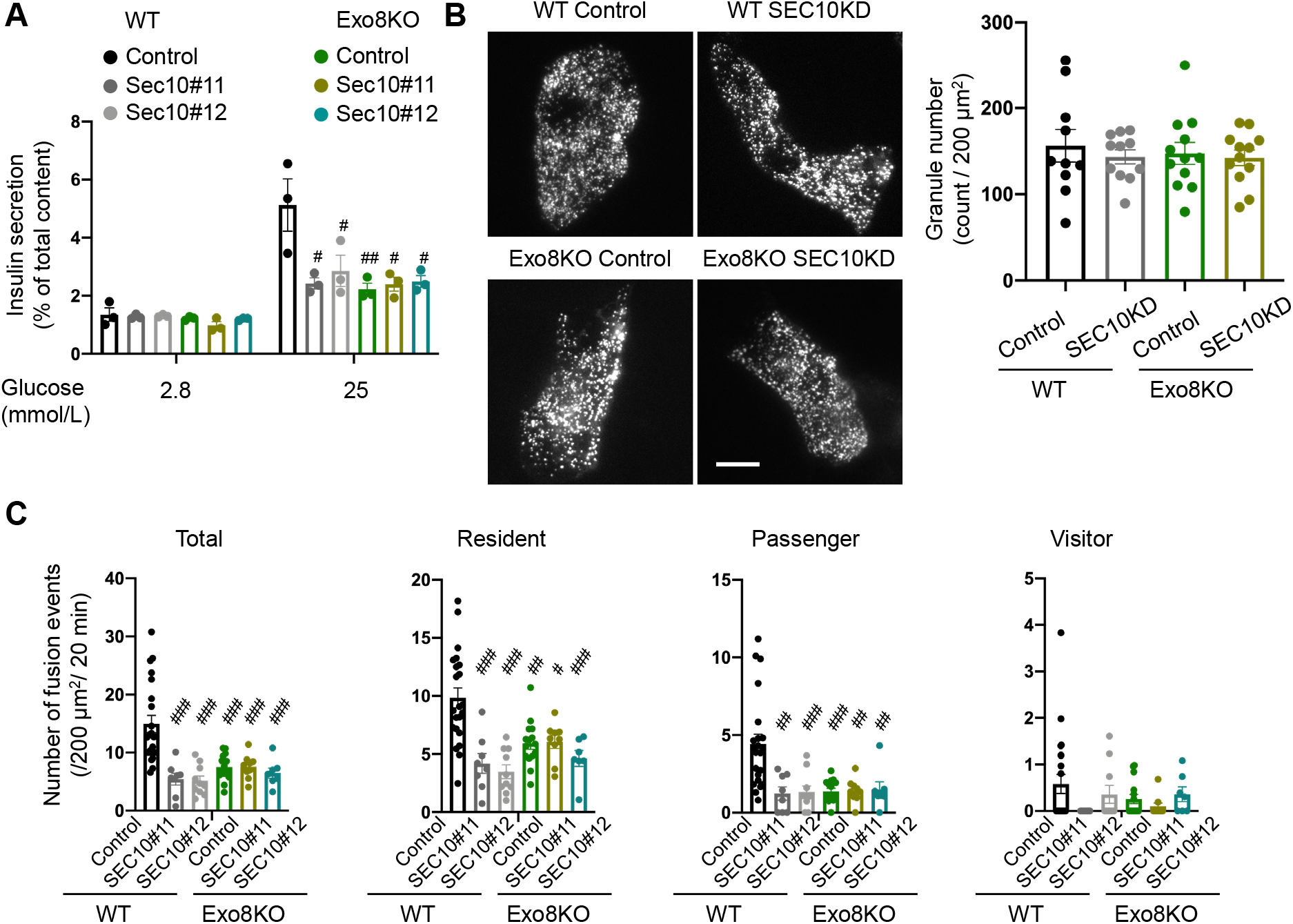
The exocyst affects insulin granule exocytosis only in the presence of exophilin-8. WT or Exo8KO mouse islet cells were twice transfected with control siRNA or siRNA against SEC10 #11 or #12, as shown in Figure 3—figure supplement 2A, and were plated in a 24-well plate (**A**) or glass base dish (**B, C**). **A**: The transfected monolayer cells (*n* = 3 from 3 mice each) were incubated for 1 h in KRB buffer containing 2.8 mmol/L glucose, and were then stimulated for 1 h in the same buffer or buffer containing 25 mmol/L glucose. The ratios of insulin secreted in the media to that left in the cell lysates are shown. **B**: The control or SEC10 knockdown (KD) cells (control siRNA-treated WT cells, *n* = 10 from 3 mice; SEC10 siRNA#11 or #12-treated WT cells, *n* = 11 from 3 mice; control siRNA-treated Exo8KO cells, *n* = 12 from 3 mice; SEC10 siRNA#11 or #12-treated Exo8KO cells, *n* = 12 from 3 mice) were infected with adenovirus encoding insulin-EGFP and were observed by TIRF microscopy (left). Numbers of visible granules were manually counted (right). Bar, 10 μm. **C**: The transfected cells (control siRNA-treated WT cells, *n* = 22 from 3 mice; SEC10 siRNA#11-treated WT cells, *n* = 8 from 3 mice; SEC10 siRNA#12-treated WT cells, *n* = 10 from 3 mice; control siRNA-treated Exo8KO cells, *n* = 15 from 3 mice; SEC10 siRNA#11-treated Exo8KO cells, *n* = 9 from 3 mice; SEC10 siRNA#12-treated Exo8KO cells, *n* = 7 from 3 mice) were infected with adenovirus encoding insulin-EGFP, and fusion events in response to 25 mmol/L glucose for 20 min were counted and categorized under TIRF microscopy as described in Figure 1D. # *P* < 0.05, ## *P* < 0.01, ### *P* < 0.001 by oneway ANOVA versus control siRNA-treated WT cells.

### In the absence of granuphilin, exophilin-8 only promotes the exocytosis of granules recruited from the cell interior to the plasma membrane after stimulation

As shown in Figure 1D, exophilin-8 deficiency also affects the resident type exocytosis. Another effector, granuphilin, is also thought to be deeply involved in this type of exocytosis, because beta cells lacking granuphilin lose almost all granules directly attached to the plasma membrane in their electron micrographs (Gomi et al., 2005). Despite this docking defect, these cells display a marked increase in granule exocytosis, possibly because granuphilin interacts with and stabilizes syntaxins in a fusion-incompetent, closed form (Torii et al., 2002). To explore the functional relationship between the two effectors in this type of exocytosis, we generated granuphilin/exophilin-8 double-knockout (GE8DKO) mice. GE8DKO cells exhibited an intermediate level of GSIS between the increased level in granuphilin-knockout (GrphKO) cells and the decreased level in Exo8KO cells (Figure 5A). These findings indicate that a significant number of granules fuse efficiently without exophilin-8 and granuphilin, thus without prior capture in the actin cortex and stable docking to the plasma membrane. TIRF microscopy of these cells expressing insulin-EGFP revealed that, although the number of visible granules was markedly decreased in GrphKO cells compared with WT cells as expected, it was not further decreased in GE8DKO cells compared with GrphKO cells (Figure 5B). The number of fusion events during glucose stimulation under TIRF microscopy was correlated with the amount of insulin released in the medium in each mutant cell (Figure 5C). All the types of exocytosis were increased in GrphKO cells, suggesting that the absence of granuphilin make granules get easier access to the fusion-competent machinery on the plasma membrane. Simultaneous absence of exophilin-8 induced strikingly differential effects on each type of exocytosis in GE8DKO cells: the increases in the passenger and visitor types were erased to the level found in Exo8KO cells, whereas the increase in the resident type was not affected at all. The former finding is consistent with the view that the passenger and visitor types of exocytosis are derived from granules captured in the actin cortex by exophilin-8. However, the latter finding indicate that at least in the absence of granuphilin, exophilin-8 is dispensable for the exocytosis from granules already residing beneath the plasma membrane before stimulation.

**Figure 5.**
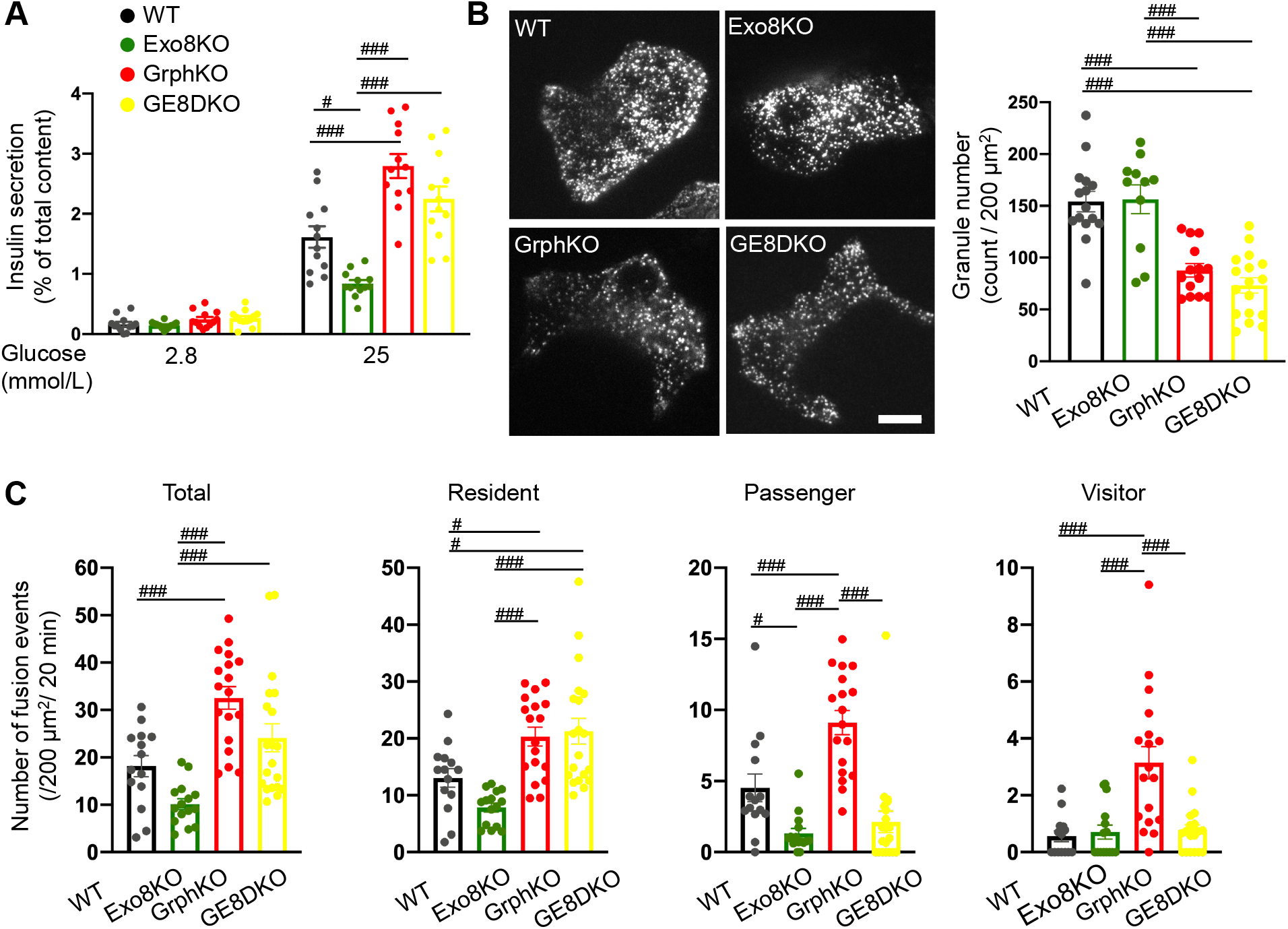
Granuphilin deficiency increases exophilin-8-independent, resident type exocytosis and exophilin-8-dependent, passenger and visitor types of exocytosis. **A**: Islets isolated from WT, Exo8KO, GrphKO, and GE8DKO mice at 12-17 weeks of ages were preincubated in LG KRB butter for 1 h. They were then incubated in another LG buffer for 30 min followed by HG buffer for 30 min. The ratios of insulin secreted in the media of that left in the cell lysates are shown as in Figure 1C (*n* = 12 from 4 mice each). **B**: A monolayer of islet cells from the above four kinds of mice were infected with adenovirus encoding insulin-EGFP and were observed by TIRF microscopy (left). Numbers of visible granules were manually counted for WT (*n* = 15 cells from 4 mice), Exo8KO (*n* = 11 cells from 3 mice), GrphKO (*n* = 14 cells from 4 mice), and GE8DKO (*n* = 17 cells from 4 mice) cells (right). Bar, 10 μm. **C:** Fusion events in response to 25 mmol/L glucose for 20 min were counted and categorized as described in Figure 1D (WT: *n* = 14 cells from 3 mice; Exo8KO: *n* = 15 cells from 3 mice; GrphKO: *n* = 18 cells from 3 mice; GE8DKO: *n* = 20 cells from 3 mice). Note that the increases in the passenger and visitor types in GrphKO cells are erased by simultaneous absence of exophilin-8 in GE8DKO cells, whereas the increase in the resident type is not affected at all. # *P* < 0.05, ### *P* < 0.001 by one-way ANOVA.

### In the presence of granuphilin, exophilin-8 also promotes the exocytosis residing beneath the plasma membrane before stimulation

However, the above findings may be a little odd, because exophilin-8 deficiency markedly affects the resident type exocytosis in the presence of granuphilin (Figure 1D, 5C). We previously showed in WT cells that the resident type exocytosis is heterogeneously derived from granuphilin-positive, immobile granules and granuphilin-negative, mobile granules (Mizuno et al., 2016). To directly assess the influence of exophilin-8 deficiency on granuphilin-mediated, docked granule exocytosis, we expressed Kusabira Orange-1 (KuO)-fused granuphilin in GrphKO and GE8DKO cells (Figure 6—figure supplement 1A), because exogenous granuphilin expressed in the presence of endogenous one abnormally accumulates insulin granules beneath the plasma membrane and severely impairs their exocytosis (Mizuno et al., 2016; Torii et al., 2004; Torii et al., 2002). Under TIRF microscopy, the number of granuphilin-positive granules as well as that of total granules were not different between the two cells. We confirmed similar numbers of these visible granules in WT and Exo8KO cells by immunostaining endogenous granuphilin and insulin (Figure 6—figure supplement 1B), suggesting that the expression levels of KuO-granuphilin in GrphKO and GE8DKO cells are properly adjusted to mimic WT and Exo8KO cells, respectively. Consistently, the number of total fusion events as well as that of the passenger type exocytosis in GE8DKO cells expressing KuO-granuphilin (mimic Exo8KO cells) were markedly decreased compared with that in GrphKO cells expressing KuO-granuphilin (mimic WT cells; Figure 6A), as found in Exo8KO cells compared with WT cells (Figure 1D, 5C). Furthermore, the number of the resident type exocytosis from granuphilin-positive granules was strongly inhibited in mimic Exo8KO cells compared with mimic WT cells. The fusion probability of granuphilin-positive granules, in which the number of fusion events is normalized by the number of visible granules, was also strongly decreased (Figure 6B). These findings indicate that exophilin-8 promotes the exocytosis of granules molecularly and stably tethered to the plasma membrane by granuphilin.

**Figure 6.**
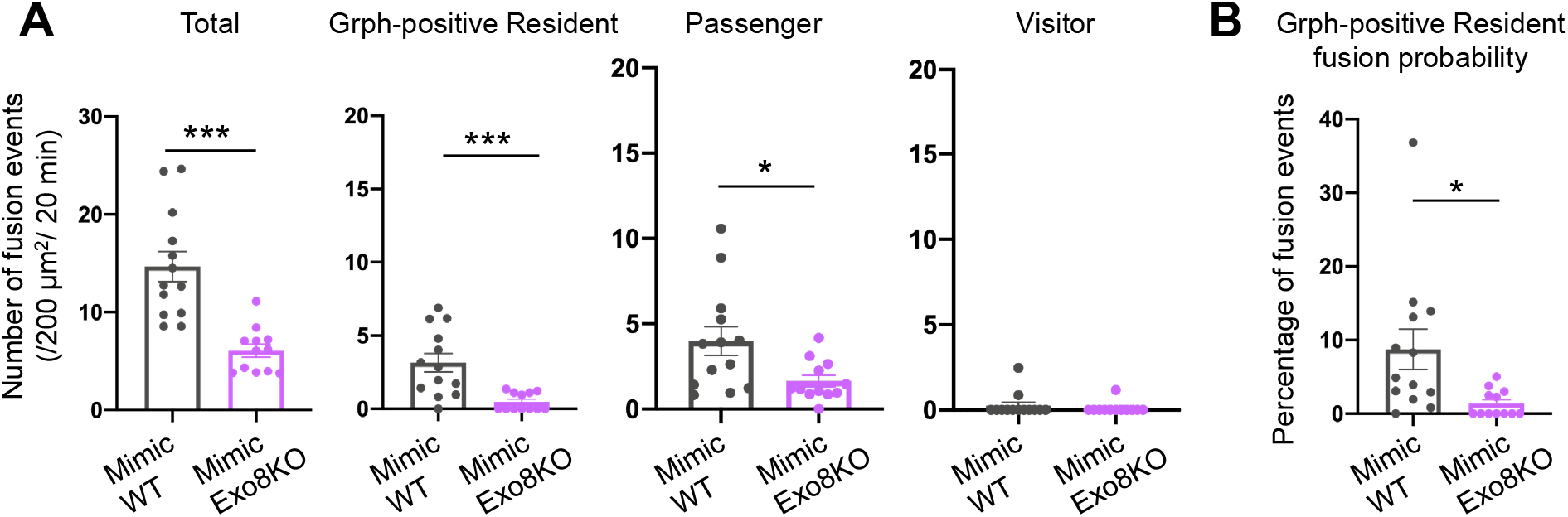
Exophilin-8 deficiency strongly inhibits the resident type exocytosis from granuphilin-positive, stably docked granules. A monolayer of GrphKO (*n* = 13) and GE8DKO (*n* = 12) islet cells from 3 mice each were infected by adenoviruses encoding insulin-EGFP and KuO-granuphilin to mimic WT and Exo8KO cells, respectively, as described in Figure 6—figure supplement 1A. Insulin granule fusion events in response to 25 mmol/L glucose for 20 min were counted and categorized under TIRF microscopy as in Figure 5C, except that granuphilin-positive granules were distinguished. There were no granuphilin-positive granules showing either passenger or visitor type exocytosis. The fusion probability of granuphilin-positive granules is shown as the percentage of those granules displaying the resident type exocytosis (**B**). * *P* < 0.05, *** *P* < 0.001 by Student *t* test.

## Discussion

In the present study, using distinct prefusion behaviors observed by TIRF microscopy as markers of differential exocytic routes, we investigated the functional hierarchy among different Rab27 effectors in pancreatic beta cells lacking one or two of them. As a result, we first present evidence that exophilin-8 functions upstream of melanophilin to drive the passenger type exocytosis. This type of exocytosis appears to be derived from granules within the actin network, because exophilin-8 is essential for granule accumulation in the actin cortex and both effectors function via interactions with myosin-VIIa and myosin-Va, respectively, in beta cells (Fan et al., 2017; Wang et al., 2020). We also show that these two effectors are linked by the exocyst protein complex. Although this evolutionarily conserved protein complex is known to function in constitutive exocytosis, it has not been established whether it plays a universal role in regulated exocytosis. For example, *Drosophila* with mutation in SEC5 displays a defect in neurite outgrowth but not in neurotransmitter secretion (Murthy et al., 2003). In mammalian cells, the exocyst complex is assembled after formation of two separate subcomplexes (Ahmed et al., 2018). The holo-exocyst appears to connect exophilin-8 and melanophilin, because both subcomplex components commonly exist in the immunoprecipitates of two effectors in beta cells. We further show that exophilin-8 and melanophilin associates via different subcomplex components, SEC8 and EXO70, respectively, and that disruption of one subcomplex results in dissociation between the two effectors. To our knowledge, this is the first example showing that different effectors toward the same Rab form a complex in cells, which corroborates the previous suggestion that the continuous presence of GTP-bound Rab27 and multiple effectors on the same granules may smooth the transition between consecutive intermediate exocytic processes (Izumi, 2021). In yeast, the exocyst interacts with Sec4, a member of the Rab protein on secretory vesicles, via exocyst component Sec15 (Guo et al., 1999). Further, Sec4 and Sec15 directly bind Myo2, the yeast myosin-V, for secretory vesicle transport (Jin et al., 2011). The exocyst components also interact with the SNARE fusion machinery: Sec6 with Snc2 (VAMP/synaptobrevin family in mammals) (Shen et al., 2013) and Sec9 (SNAP-25 family in mammals) (Dubuke et al., 2015), and Sec3 with Sso2 (syntaxin family in mammals) (Yue et al., 2017). Although proteins corresponding to Rab27 effectors are missing in yeast, they appear to be involved in similar interactions in mammalian cells. In fact, melanophilin interacts with Rab27a, myosin-Va, and syntaxin-4 in beta cells (Wang et al., 2020).

We then show that exophilin-8 knockout and SEC10 knockdown exhibit very similar defects in granule exocytosis, indicating that both function together. The sole difference is that, although exophilin-8 deficiency erases the granule accumulation in the actin cortex, exocyst deficiency does not, which suggests that the exocyst functions downstream of exophilin-8. Consistent with this view, we could not find any effects of SEC10 knockdown in Exo8KO cells. However, the exocyst should function upstream of melanophilin, because its deficiency not only decreases the passenger type exocytosis, but also markedly reduces the resident type exocytosis that should be derived from granules residing within a 200 nm from the plasma membrane before stimulation. Surprisingly, however, SEC10 knockdown does not affect the number of such granules visualized by TIRF microscopy, although the exocyst is known to physically tether secretory vesicles to the plasma membrane prior to membrane fusion in a broad range of cells (Wu and Guo, 2015). Although this may be due to partial downregulation by SEC10 siRNA in our experiments, it should be noted that granule exocytosis is still markedly reduced under this condition. The absence of exophilin-8 has no effect on granule residence in the evanescent area either, which suggests that granules can get access to the plasma membrane without prior capture in the actin cortex. In the pheochromocytoma cell line, PC12, nascent granules generated are transported in a microtubule-dependent manner within a few seconds to the cell periphery (Rudolf et al., 2001). In skin melanocytes, melanosomes are dispersed throughout the cytoplasm using myosin-Va motor along dynamic actin tracks assembled by the SPIRE actin nucleator (Alzahofi et al., 2020). Thus, insulin granules may also be eventually transported close to the plasma membrane using these routes. They are then thought to be stably docked to the plasma membrane by another Rab27 effector, granuphilin (Gomi et al., 2005; Mizuno et al., 2016).

We finally show that exophilin-8 deficiency has differential effects on the resident type exocytosis: a strong inhibition in the presence of granuphilin and no inhibition despite its increase in the absence of granuphilin. Considering that exophilin-8 deficiency has no effect on granule numbers residing beneath the plasma membrane whether granuphilin is present or not, and that it almost completely inhibits the exocytosis of granuphilin-positive, stably docked granules, exophilin-8 should have a role in something other than granule location. Granuphilin-mediated, stably docked granules are thought to require priming machinery for fusion, such as Munc13, that converts a granuphilin-associated, closed form of syntaxin to a fusion-competent, open form (Mizuno and Izumi, 2022). Exophilin-8 can contribute to this process, because it is associated with RIM-BP2, RIM, and Munc13 (Fan et al., 2017), which are known to have such a priming role in synaptic vesicle exocytosis (Brockmann et al., 2019; Brockmann et al., 2020). In fact, the exophilin-8 mutant that loses the binding activity to RIM-BP2 fails to rescue the decreased insulin secretion in exophilin-8-deficient cells (Fan et al., 2017).

In summary, our findings present the first diagram for the functional hierarchy among different Rab27 effectors expressed in the same cell (Figure 7). It is now evident that there are multiple redundant paths and rate-limiting processes toward the final fusion step in granule exocytosis, which should secure the robustness of mechanism secreting vital molecules, such as insulin.

**Figure 7.**
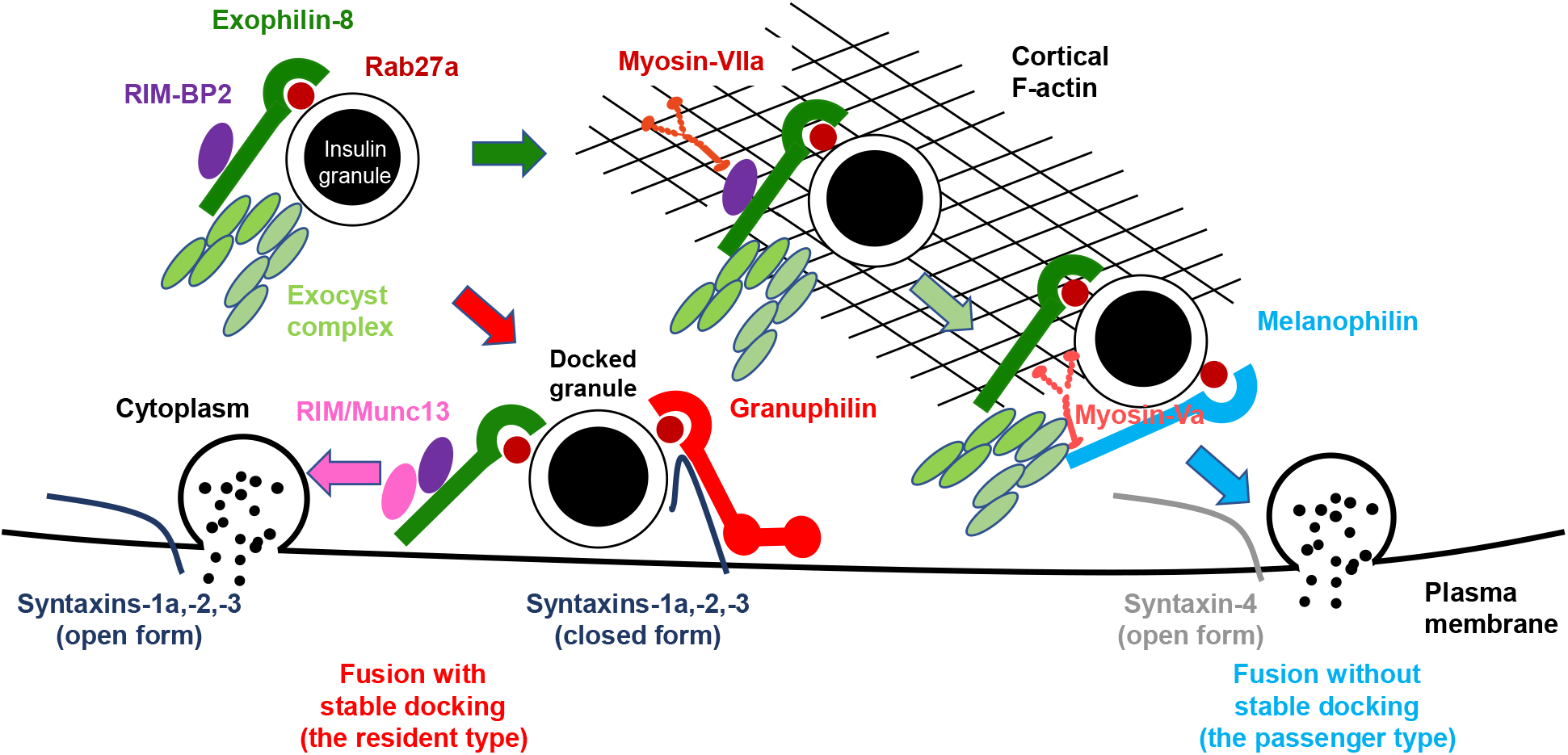
A schematic model for the functional relationship among different Rab27 effectors, exophilin-8, melanophilin, and granuphilin, in insulin granule exocytosis. A part of granules in the cytoplasm are captured within the actin cortex by exophilin-8 that interacts with RIM-BP2 and myosin-VIIa (dark green arrow). The exocyst associates exophilin-8 to melanophilin (light green arrow), which mediates the passenger type exocytosis via interactions with myosin-Va and the fusion-competent, open form of syntaxin-4 (light blue arrow). Other granules in the cytoplasm are transported to the cell periphery, perhaps along microtubule or actin tracks, in an exophilin-8-independent manner, and are then stably docked to the plasma membrane by granuphilin that prevents spontaneous fusion by interacting with the fusion-incompetent, closed form of syntaxin-1a, 2, and 3 (red arrow). Exophilin-8 and the exocyst are also involved in the resident type exocytosis, possibly by carrying the exophilin-8-associated RIM-BP2, RIM, and Munc13 that convert granuphilin-associated syntaxins to the open form (pink arrow).

## Materials and methods

### Mice and pancreatic islet cell preparation

Animal experiments were performed according to the rules and regulations of the Animal Care and Experimental Committees of Gunma University (permit number: 22-010; Maebashi, Japan). Only male mice and their tissues and cells were phenotypically characterized in this study. *Leaden* (C57J/L) mice with nonfunctional mutation of the gene encoding melanophilin (Matesic et al., 2001) were purchased from The Jackson Laboratory (Strain #:000668, RRID: IMSR_JAX:000668), and were backcrossed with C57BL/6N mice ten times to generate MlphKO mice. Exo8KO mice in the genetic background of C57BL/6N mice were described previously (Fan et al., 2017). ME8DKO mice were obtained by mating Exo8KO mice with MlphKO mice described above. GrphKO mice in the genetic background of C3H/He mice were described previously (Gomi et al., 2005). The male Exo8KO mice were mated with the female GrphKO mice. Because the granuphilin and exophilin-8 genes are on mouse X and 9 chromosomes, respectively, the resultant F1 generation is either male (*Grph^-/Y^, Exo8^+/-^*) or female (*Grph^+/-^, Exo8^+/-^*) mice. By crossing these F1 mice, GE8DKO mice, as well as WT, GrphKO, and Exo8KO mice, were generated in the F2 generation, and used for experiments. Although those F2 mice have a mixture of C57BL/6N and C3H/He genomes, we expected that significant phenotypic changes due to the loss of Rab27 effectors may be preserved beyond the noises due to randomly distributed differences in the genome. In fact, we found similar differences in exocytic profiles between WT and Exo8KO cells both in the C57BL/6N background (Figure 1) and in the mixture of C57BL/6N and C3H/He background (Figure 5). Furthermore, GrphKO cells in the mixture of C57BL/6N and C3H/He background show changes in granule localization and exocytosis (Figure 5), consistent with the reported phenotypes of GrphKO cells in the C3H/He background (Gomi et al., 2005). Pancreatic islet isolation and dissociation into monolayer cells and insulin secretion assays were performed as described previously (Gomi et al., 2005; Wang et al., 2020). Briefly, islets were isolated from cervically dislocated mice by pancreatic duct injection of collagenase solution, and size-matched five islets were cultured overnight in a 24-well plate. Monolayer islet cells were prepared by incubation with trypsin-EDTA solution, and were cultured for further 2 days. Insulin released from isolated islets or monolayer cells was measured by an AlphaLISA insulin kit (PerkinElmer) or an Insulin high range kit (Cisbio).

### DNA manipulation

Human cDNAs of SEC3, SEC5, SEC6, SEC8, SEC10, SEC15, EXO70, and EXO84 in the pEGFP-C3 vector were gifts from Dr. Channing J. Der (Addgene plasmid # 53755-53762; http://n2t.net/addgene: 53755-53762; RRID: Addgene_53755-53762) (Martin et al., 2014). Adenoviruses encoding insulin-EGFP and KuO-granuphilin were described previously (Kasai et al., 2008; Mizuno et al., 2016). Hemagglutinin (HA)-, FLAG-, MEF-, One-STrEP-FLAG (OSF), and mCherry-tagged exophilin-8 and melanophilin were made previously (Fan et al., 2017; Wang et al., 2020). To express exogenous protein, HEK293A cells were transfected with the plasmids using Lipofectamine 3000 reagent (Invitrogen), whereas MIN6 and primary pancreatic beta cells were infected with adenoviruses.

### Cell lines, antibodies, and immunoprocedures

MIN6 (RRID: CVCL_0431) and HEK293A (RRID: CVCL_6910) cells were listed by NCBI Biosample (SAMEA4168040 and SAMEA4146837), respectively. Guinea pig anti-insulin serum is a gift from H. Kobayashi (Gunma University). Rabbit polyclonal anti-exophilin-8 (αExo8N) and anti-granuphilin (αGrp-N) antibodies are described previously (Fan et al., 2017; Yi et al., 2002). mCherry nanobody is a gift from Drs. Y. Katoh and K. Nakayama (Kyoto University) (Katoh et al., 2016). The sources of commercially available antibodies and their concentrations used are listed in Supplementary Table. Cells were lysed by lysis buffer (50 mmol/L Tris-HCl, pH 7.5, 150 mmol/L NaCl, 10% (w/v) glycerol, 100 mmol/L NaF, 10 mmol/L ethylene glycol tetraacetic acid, 1 mmol/L Na_3_VO_4_, 1% Triton X-100, 5 μmol/L ZnCl_2_, 1 m mmol/L phenylmethylsulfonyl fluoride, and complete protease inhibitor cocktail (Roche)). Immunoblotting and immunoprecipitation were performed as described previously (Matsunaga et al., 2017; Wang et al., 2020). VIP assay using the nanobody was performed as described previously (Katoh et al., 2015). Briefly, HEK293A cells on 10 cm dish were transfected with pEGFP-Sec and either pmCherry-Melanophilin or pmCherry-Exophilin-8 by Lipofectamine 3000. After 48 h, cells were lysed by 1ml of lysis buffer, and the cell lysate was centrifuged at 10,000× g for 10 min, and the supernatant was subjected to immunoprecipitation with 5 μl gel volume of mCherry nanobody-bound glutathione Sepharose. The beads were washed three times with lysis buffer and transferred to 35-mm glass base dish (Glass φ12, IWAKI). Green and red fluorescence of beads were observed by confocal laser scanning microscopy. Acquisition of images were performed under fixed conditions. For immunofluorescence procedures, monolayer primary beta cells seeded at 3-5 × 10^4^ cells on poly-L-lysine-coated glass base dish were fixed by 4% paraformaldehyde for 30 min at room temperature. The cells were washed by phosphate buffered saline (PBS) and permeabilized by PBS containing 0.1% Triton X-100 and 50 mmol/L NH4Cl for 30 min. After blocking with 1% bovine serum albumin in PBS for 30 min, the cells were immunostained and observed under confocal microscopy with a 100× oil immersion objective lens (1.49 NA). Quantification of immunoblot signals was performed using ImageQuant TL software (Cytiva). Quantification of colocalization between two proteins was performed using NIS Element Viewer software (Nikon).

### TIRF microscopy

TIRF microscopy was performed as described previously (Wang et al., 2020). Briefly, monolayer islet cells on glass base dish were infected with adenovirus encoding insulin-EGFP. Two days after, the cells were preincubated for 30 min in 2.8 mmol/L low glucose (LG)-containing Krebs-Ringer bicarbonate (KRB) buffer at 37°C. They were then incubated in new LG buffer for 20 min or 25 mmol/L high glucose (HG)-containing buffer for 20 min. TIRF microscopy was performed on 100× oil immersion objective lens (1.49 NA). The penetration depth of the evanescent field was 100 nm. Images were acquired at 103-ms intervals. Fusion events with a flash followed by diffusion of EGFP signals were manually selected and assigned to one of three types: residents, which are visible over 10 s before fusion; visitors, which have become visible within 10 s before fusion; and passengers, which are not visible before fusion. In case of coinfection with adenovirus encoding KuO-granuphilin, sequential multi-color TIRF microscopy was performed as described previously (Mizuno et al., 2016). Briefly, EGFP was excited using a 488-nm solid-state laser, whereas KuO was excited using a 561-nm laser. Excitation illumination was synchronously delivered from an acousto-optic tunable filter-controlled laser launch. A dualband filter set (LF488/561-A; Semrock) was applied on a light path.

### Mass spectrometry

MIN6 cells (2 × 10^7^ cells in ten 15 cm dishes) were infected with adenovirus encoding FLAG, FLAG-melanophilin, or MEF-exophilin-8. The anti-FLAG immunoprecipitates were subjected to gel electrophoresis and visualized by Oriole fluorescent gel staining (BioRad). Specific bands were excised and digested in gels with trypsin, and the resulting peptide mixtures were analyzed by a LC-MS/MS system, as described previously (Matsunaga et al., 2017). All MS/MS spectra were separated against the Mus musculus (mouse) proteome data set (UP000000589) at the Uniplot using Protein pilot software (SCIEX).

### Silencing of SEC10 in mouse pancreatic islet cells

On-Target plus Set of 4 siRNA (J-047583-11 and −12) against mouse Exoc5 (catalog no. 105504) and On-Target plus non-targeting pool siRNA were purchased from Horizon Discovery Ltd. Mouse pancreatic islet cells suspended in 1 × 10^5^ cells/280 μl were transfected with 100 nmol/L siRNA using Lipofectamine RNAiMAX reagent (Invitrogen). After plated on a 24-well plate or glass base dish for 48 h, the cells were transfected with the same siRNA for the second time. After another 24 h, the cells were subjected to immunoblotting analyses, immunofluorescent staining, insulin secretion assays, or TIRF microscopy after infection with adenovirus encoding insulin-EGFP.

### Statistical analysis

All quantitative data are assessed as the mean ± SEM. The *P* values were calculated using the Student *t* test or a one-way ANOVA with a Tukey multiple-comparison, using GraphPad Prism software.

### Data and resource availability

All data generated or analyzed during this study are included in the manuscript and supporting files. All noncommercially available resources generated and/or analyzed during the current study are available from the corresponding author upon reasonable request.

## Acknowledgments

The authors thank T. Nara, E. Kobayashi, and T. Ushigome for colony maintenance of mice, and S. Shigoka for preparing the manuscript. This work was supported by Japan Society for the Promotion of Science KAKENHI grants JP19H03449 to T.I., JP20K06535 to K.Mi, and JP20K15742 to H.W. It was also supported by grants from Eli Lilly Japan Donation and Teijin Pharma Donation to T.I.

---

**Figure 1—figure supplement 1.**
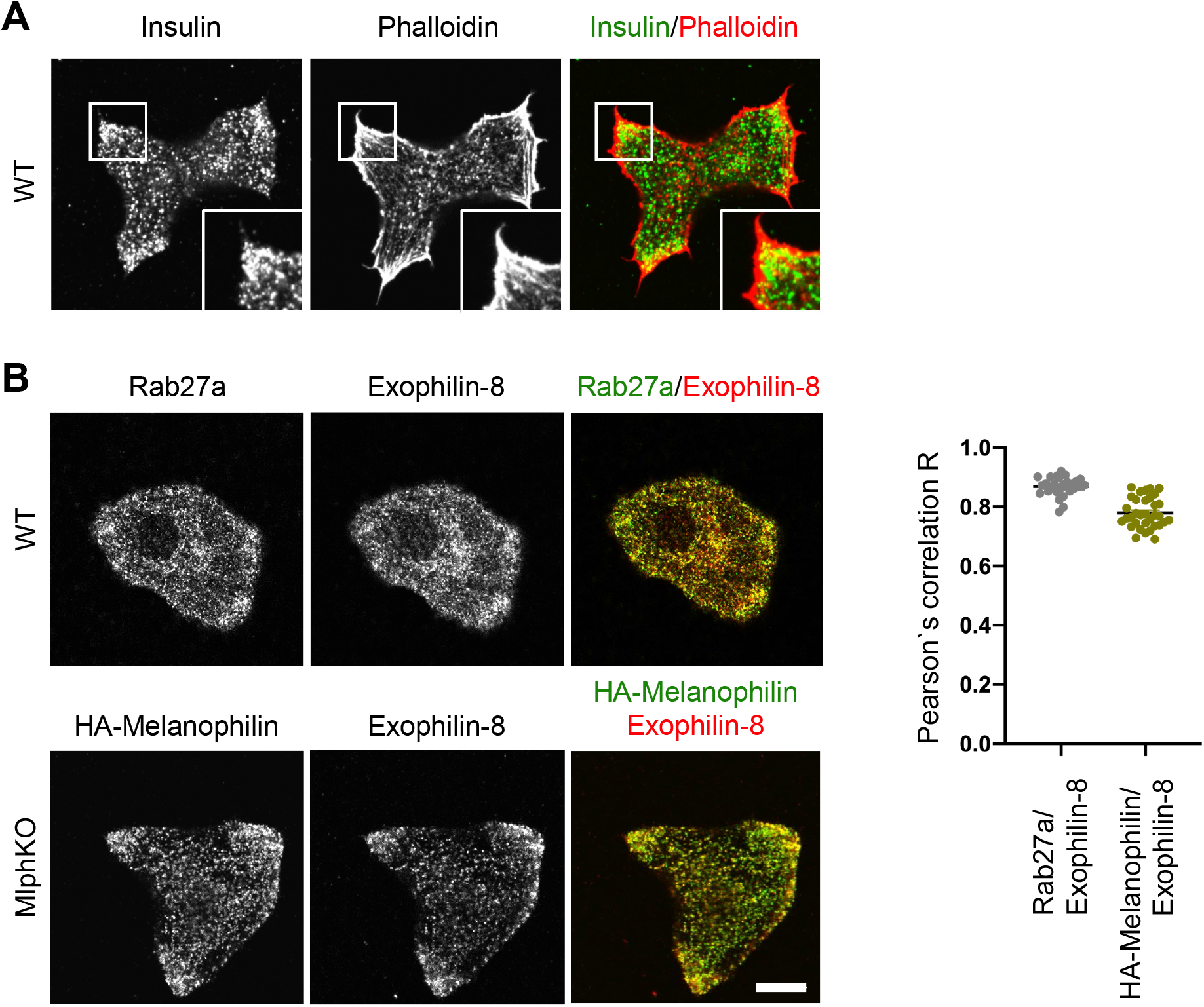
Melanophilin and exophilin-8 are colocalized on insulin granules accumulated in the actin-rich cell periphery. **A**: Primary beta cells from WT mice were double-stained with rhodamine-conjugated phalloidin and anti-insulin antibody, and were observed by confocal microscopy. **B**: WT (upper) and MlphKO beta cells expressing HA-melanophilin at the endogenous level in WT cells (lower) were coimmunostained with goat anti-exophilin-8 and either anti-Rab27a (upper) or anti-HA antibodies (lower). We investigated the intracellular distribution of HA-melanophilin expressed in MlphKO beta cells, because there is no available anti-melanophilin antibody durable for staining endogenous melanophilin, and because exogenous melanophilin expressed in the presence of endogenous one displays aberrant localization along F-actin (Wang et al., 2020). Colocalization between exophilin-8 and Rab27a in WT cells and that between exophilin-8 and HA-melanophilin in MlphKO cells were quantified by Pearson’s correlation coefficient (*n* = 30-37 from 3 mice each). Note that exophilin-8 and HA-melanophilin are accumulated and colocalized in the cell periphery. Bars, 10 μm.

**Figure 2—figure supplement 1.**
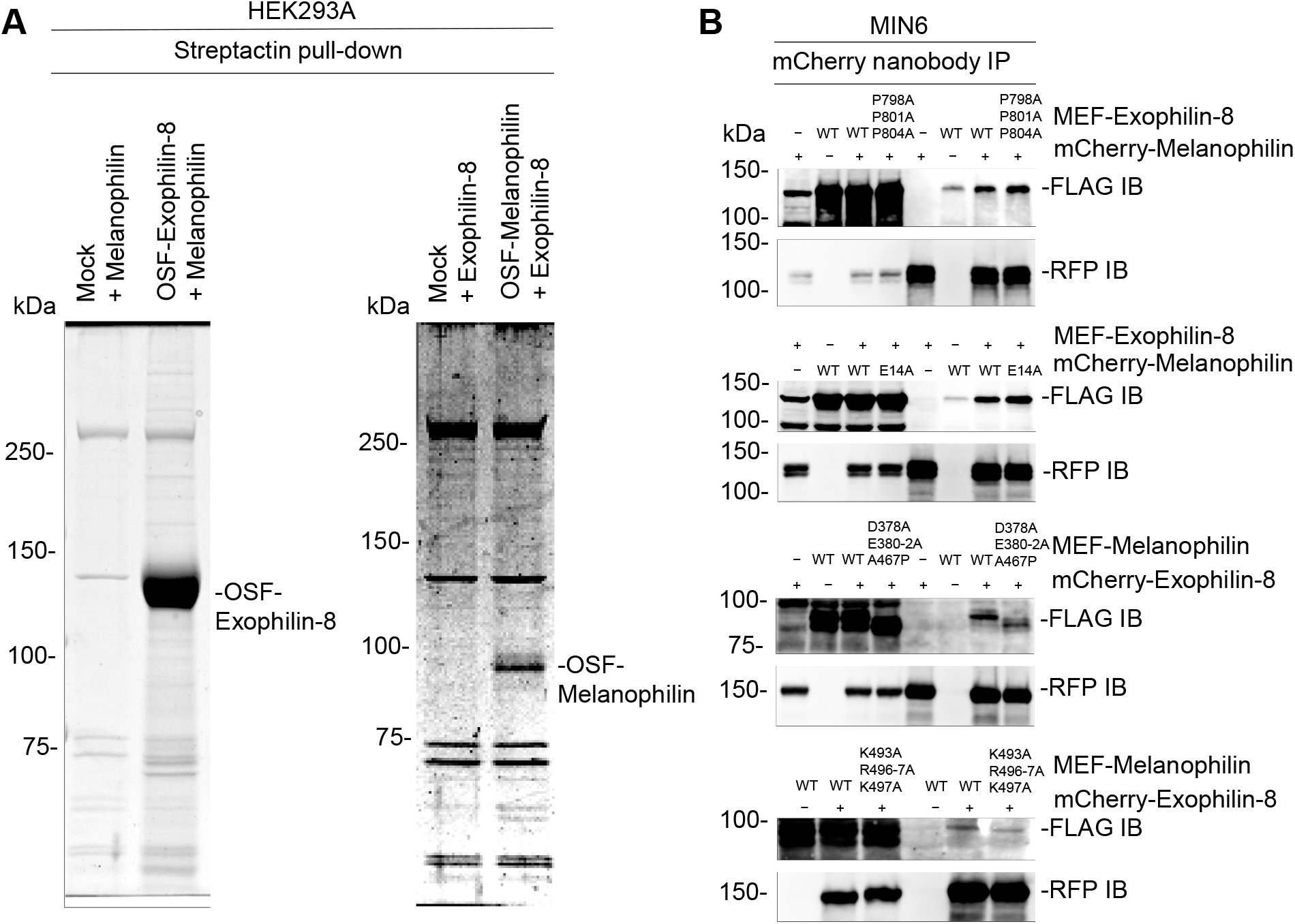
Exophilin-8 does not bind melanophilin directly or through the known interacting proteins. **A**: Melanophilin and OSF-tagged exophilin-8 (left), or exophilin-8 and OSF-melanophilin (right) were coexpressed by transfection in HEK293A cells. The cell extracts were pulled down using Strep-Tactin beads, and were subjected to gel electrophoresis followed by Coomassie Brilliant Blue gel staining. Note that exogenously expressed exophilin-8 and melanophilin do not interact in HEK293A cells. **B:** MEF-tagged, WT exophilin-8 and its P798A/P801A/P804A mutant that loses the binding activity toward RIM-BP2 were expressed with or without mCherry-melanophilin by adenoviruses in MIN6 cells (top panel). Similarly, mCherry-melanophilin WT or its E14A mutant that loses the binding activity toward Rab27a were coexpressed with or without MEF-exophilin-8 (second panel), whereas MEF-melanophilin D378A/E380A/E381A/E382A/A467P and K493A/R495A/R496A/K497A mutants that lose the binding activity toward myosin-Va and actin, respectively, were coexpressed with or without mCherry-exophilin-8 (third and bottom panels). The cell extracts underwent immunoprecipitation with anti-mCherry nanobody, and the immunoprecipitates were immunoblotted with anti-FLAG and anti-RFP antibodies. Note that any of the exophilin-8 and melanophilin mutants preserves the interaction with the other WT effector.

**Figure 2—figure supplement 2.**
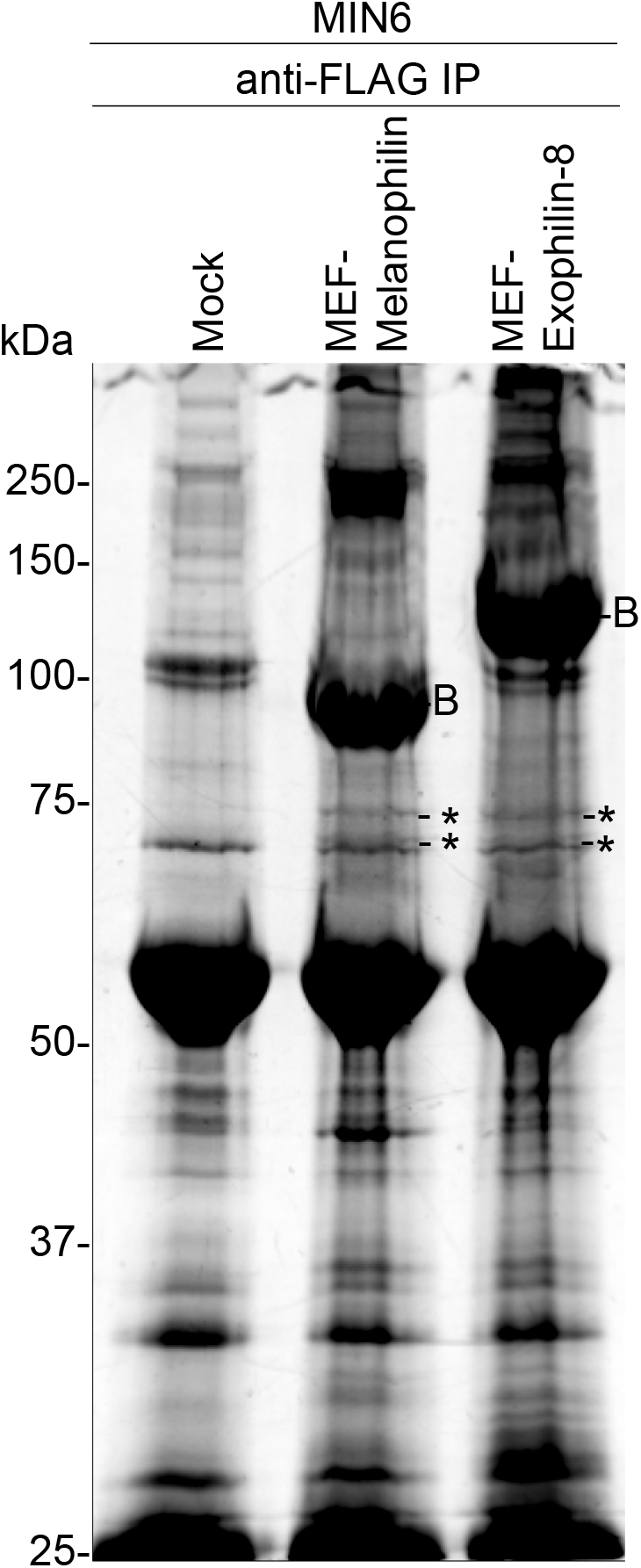
Analysis of the melanophilin and exophilin-8 protein complexes. MIN6 cells were infected by adenoviruses encoding no protein (Mock), MEF-tagged melanophilin or exophilin-8. The binding proteins were pulled down using FLAG beads, and were subjected to gel electrophoresis followed by Oriole fluorescent gel staining. The specific protein bands were cut and digested for LC-MS/MS. Asterisks and ‘B’ indicate the bands containing the exocyst complex components and bait protein, respectively.

**Figure 3—figure supplement 1.**
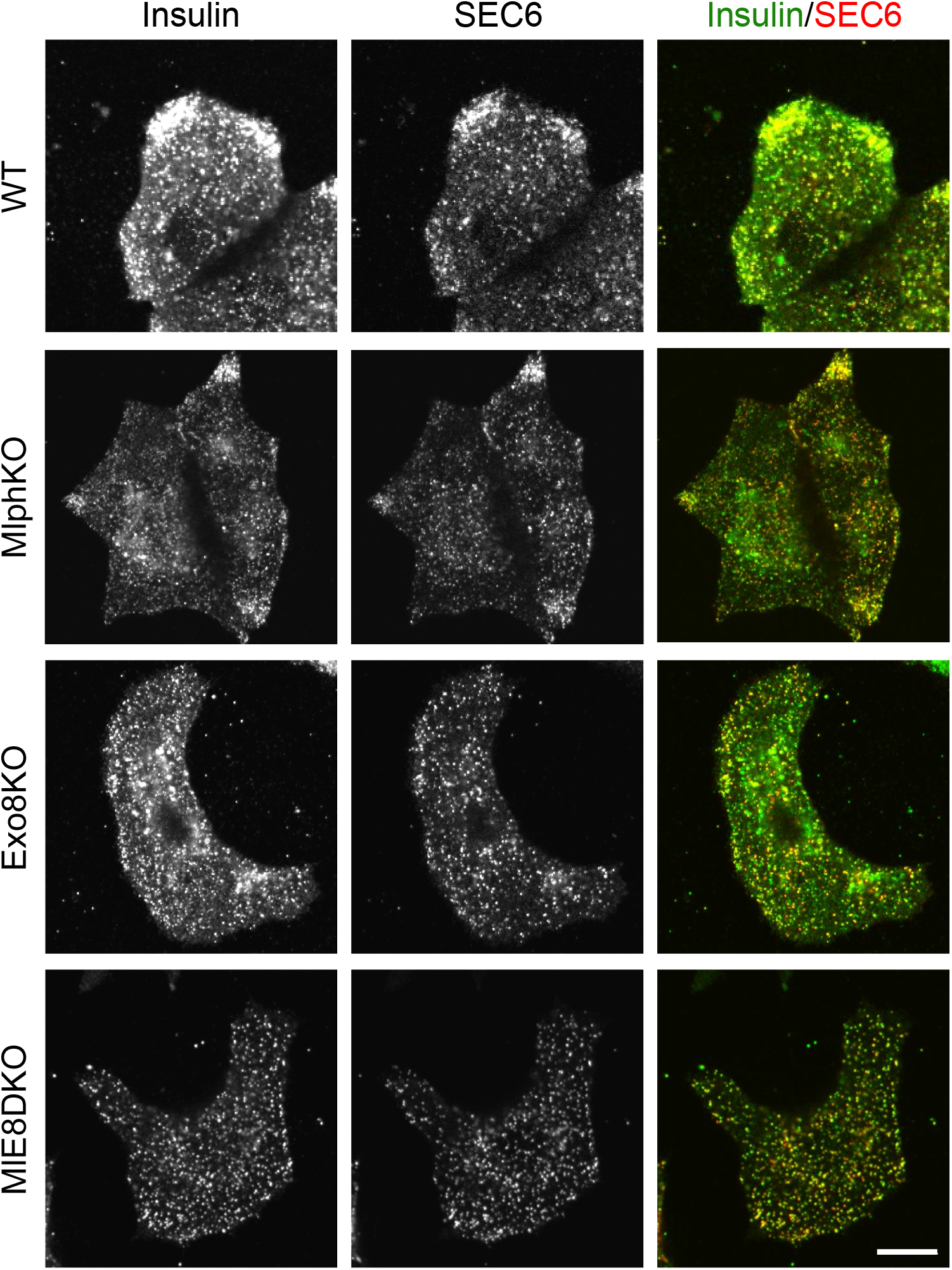
The exocyst component, SEC6, is localized on insulin granules in the absence of exophilin-8 and melanophilin. WT, MlphKO, Exo8KO, and ME8DKO beta cells were coimmunostained with anti-insulin and anti-SEC6 antibodies. Note that SEC6 is localized on insulin granules evenly in the absence of exophilin-8, although it is accumulated on granules at the cell periphery in the presence of exophilin-8. Bar, 10 μm.

**Figure 3—figure supplement 2.**
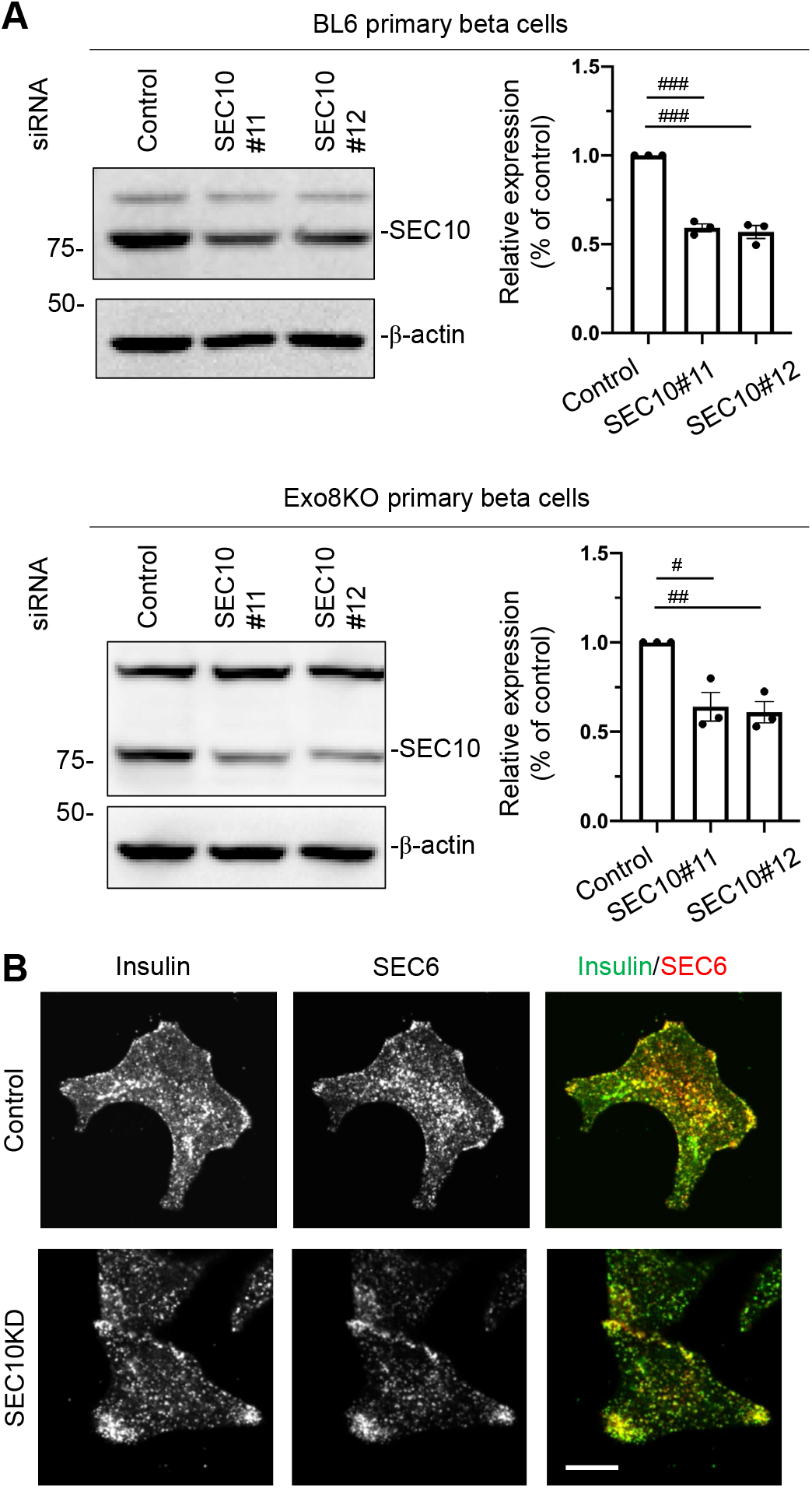
Downregulation of SEC10 in mouse pancreatic beta cells. **A**: Mouse pancreatic islet cells from WT and Exo8KO mice were suspended in 1 ×10^5^ cells/280 μl, and were transfected with 100 nmol/L control siRNA or siRNAs against SEC10 #11 or #12. After being plated on the 24-well plate for 48 h, the cells were transfected with siRNAs for the second time. After another 48 h, the cell extracts were immunoblotted with anti-SEC10 and anti-β-actin antibodies. Protein bands were quantified by densitometric analyses from three independent preparations. # *P* < 0.05, ## *P* < 0.01, ### *P* < 0.001 by one-way ANOVA. **B**: WT cells with or without SEC10 knockdown (KD) were coimmunostained with anti-insulin and anti-SEC6 antibodies, and were observed by confocal microscopy. Bar, 10 μm. **Source data 3.** Uncropped blot images of figure supplement 2A.

**Figure 6—figure supplement 1.**
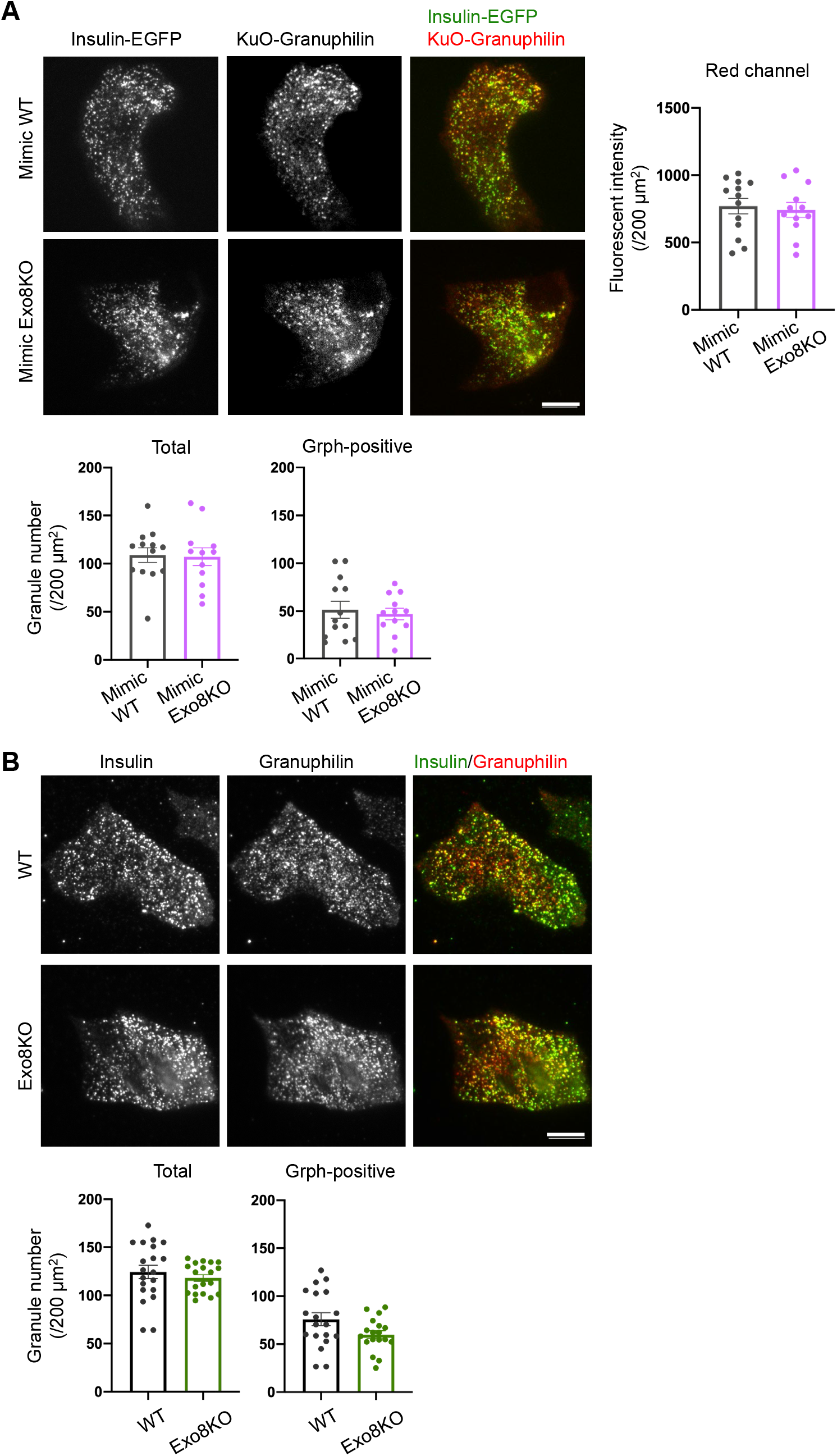
Visible insulin granules associated with granuphilin under TIRF microscopy. **A**: A monolayer of GrphKO (*n* = 13) and GE8DKO (*n* = 12) islet cells from 3 mice each were infected by adenoviruses encoding insulin-EGFP and KuO-granuphilin to mimic WT and Exo8KO cells, respectively. These cells were observed by TIRF microscopy (upper left), and the fluorescent intensities of red channel were measured using NIS Elements Viewer software to confirm the similar expression levels of KuO-granuphilin (upper right). The numbers of total and KuO-granuphilin-positive, visible granules were counted under TIRF microscopy and were compared by Student *t* test, as in Figure 5B (lower). **B**: A monolayer of WT (*n* = 19) and Exo8KO (*n* = 18) islet cells from 3 mice each were immunostained with anti-insulin and anti-granuphilin antibodies and were observed by TIRF microscopy (upper). Total and granuphilin-positive granules were counted and compared as in **A** (lower). Bars, 10 μm.

**Supplementary Table.**
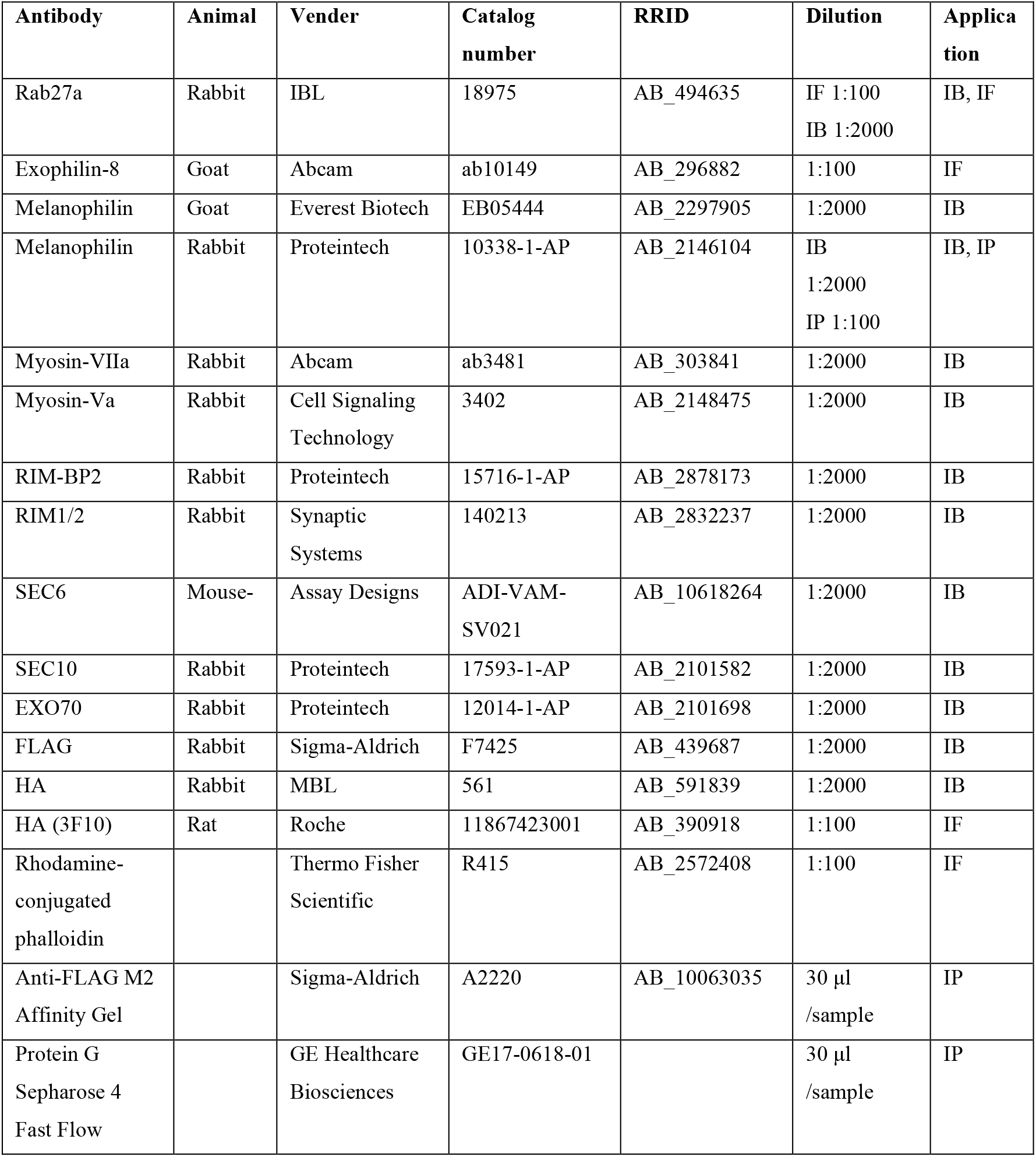
The sources of commercially available antibodies and the concentrations of the antibodies used for immunofluorescence (IF), immunoblotting (IB), or immunoprecipitation (IP)

